# Elevated Neurokinin 1-Receptor Expression in uterine products of conception is associated with first trimester Miscarriages

**DOI:** 10.1101/2020.06.27.175273

**Authors:** Riffat Mehboob, Syed Amir Gilani, Amber Hassan, Adeel Haider Tirmazi, Fridoon Jawad Ahmad, Javed Akram

## Abstract

**Background:** Miscarriage is a common complication of early pregnancy, mostly occurring in first trimester. However, the etiological factors, prognostic and diagnostic biomarkers are not well known. Neurokinin-1 Receptor (NK-1R) is a receptor of tachykinin peptide, Substance P (SP) and has a role in various pathological conditions, cancers but it’s association with miscarriages and significance as a clinicopathological parameter is not studied. Accordingly, the present study aimed to clarify the localization and expression for NK-1R in human retained products of conception. Role of NK-1R is not known in miscarriages.

**Materials and Methods:** NK-1R expression was assessed in products of conception by immunohistochemistry. Protein expression was evaluated using the nuclear labelling index (%).

**Results:** Ten human products of conception tissues were studied by immunohistochemistry to demonstrate the localization of NK-1R. The expression of NK-1R protein was high in all the cases of POCs. NK-1R expression showed no notable differences among different cases of miscarriages irrespective of the mother’s age and gestational age at which the event occurred.

**Conclusions:** Expression of NK-1R was similar in all the cases and it was intense. It shows that dysregulation of NK-1R along with its ligand Substance P might be involved in miscarriages. Our results provide fundamental data regarding this anti-NK-1R strategy. Thus, the present study recommends that SP/NK1R system might, therefore, be considered as an emerging and promising diagnostic and therapeutic strategy against miscarriages. Hence, we report for the first time, the expression and localization of NK-1R in products of conception. We suggest NK-1R antagonist in addition to the Immunoglobulins and Human chorionic gonadotropin, to diagnose and treat spontaneous miscarriages.

## Introduction

Miscarriages or spontaneous abortion in the initial stage of pregnancy is a common problem in first trimester of pregnancy [1]. Many factors are involved in it and it is a complex phenomenon(<u>27</u>). Approximately, 15% of the pregnant females miscarry spontaneously without any known cause [2]. Miscarriages are further divided into incomplete, complete, missed and anembryonic miscarriages [3]. 50% of causes are still unknown and no indicators have been identified in them. Tests to diagnose these cases include an ultrasound scanning and measurement of human chorionic gonadotropin. In this way, the patients at risk are identified. However, there is a dire need to explore potential efficient biomarkers for the diagnosis of at risk patients earlier than the onset of clinical symptoms or the unfortunate occurrence of event [4]. It will not only provide a diagnostic strategy, but also pave way for therapeutic interventions to manage such cases. For this purpose, Neurokinin-1 receptor (NK-1R) is explored in the current study to evaluate its expression and localization in the products of conception (POC) tissues after a miscarriage.

NK-1 is a receptor of Substance P (SP) protein, which is one of the peptides released from sensory nerves, causes the enhancement of cellular excitability in several human tumor cells[5]. There are two distinct conformational isoforms of NK-1R: a full-length NK-1R (NK-1RF) isoform, a truncated NK-1R (NK-1RT) isoform. Truncated NK-1R (NK-1RT) isoform lacks the terminal cytoplasmic 96-aa residues [6]. Both of these isoforms have same binding affinity for SP but different affinities for NKA. The NK-1R has a relatively long 5◻ untranslated region compared to the other TK receptors, which is preceded by a single TATAAA sequence [7].

The neurokinins are a class of peptide signaling molecules that mediate a range of central and peripheral functions including pain processing, gastrointestinal function, stress responses, like hematopoiesis[8], wound healing[9], increased vascular permeability, neurogenic inflammation, leukocyte infiltration, cell survival and anxiety[10]. It is also involved in carcinogenesis as reported in many studies and lead to metastasis [11–14]. SP has a variety of physiological functions in humans, particularly, nervous, immune and cardiovascular systems [15]. Mainly, SP is a braingut hormone, and its receptor is present in brain regions but also in peripheral tissues. SP binds to NK-1R to carryout transmission of pain, secretions from paracrine and endocrine system, vasodilation and proliferations of cells [16, 17]. It is a neuromodulator, neurotransmitter as well as a neurohormone.

Tachykinins (TKs) possess a wide spread distribution in central & peripheral nervous system that is undoubtedly a major source of these peptides. However TKs have also a limited distribution in non-neural structures represented by the irregular and sparse localizations in which they display known and unknown functions. In neuronal cells, the active TKs act as neurotransmitters/neuromodulators. In the non-neuronal cell, as autocrine, paracrine and endocrine regulators.

SP brings all of its cellular activities after binding to the G-protein coupled receptor (GPCR) NK-1R (Figure 1). NK-1R has 407 amino acids and is encoded by TACR-1 gene. It is located on cell surface [18]. SP binds to heterotrimeric GPCR, with a preference for Gs and Gq like all TK receptors. Binding of receptor to Gs stimulates adenylyl cyclase and cyclic AMP production whereas binding of receptor to Gq stimulate the phosphatidyl-inositol casade [19]. Upon binding of SP to NK-1R, a signal transduction cascade is initiated by internalization of SP-NK-1R complex [20] that activates the Phospholipase C (PLC).

**Figure 1:**
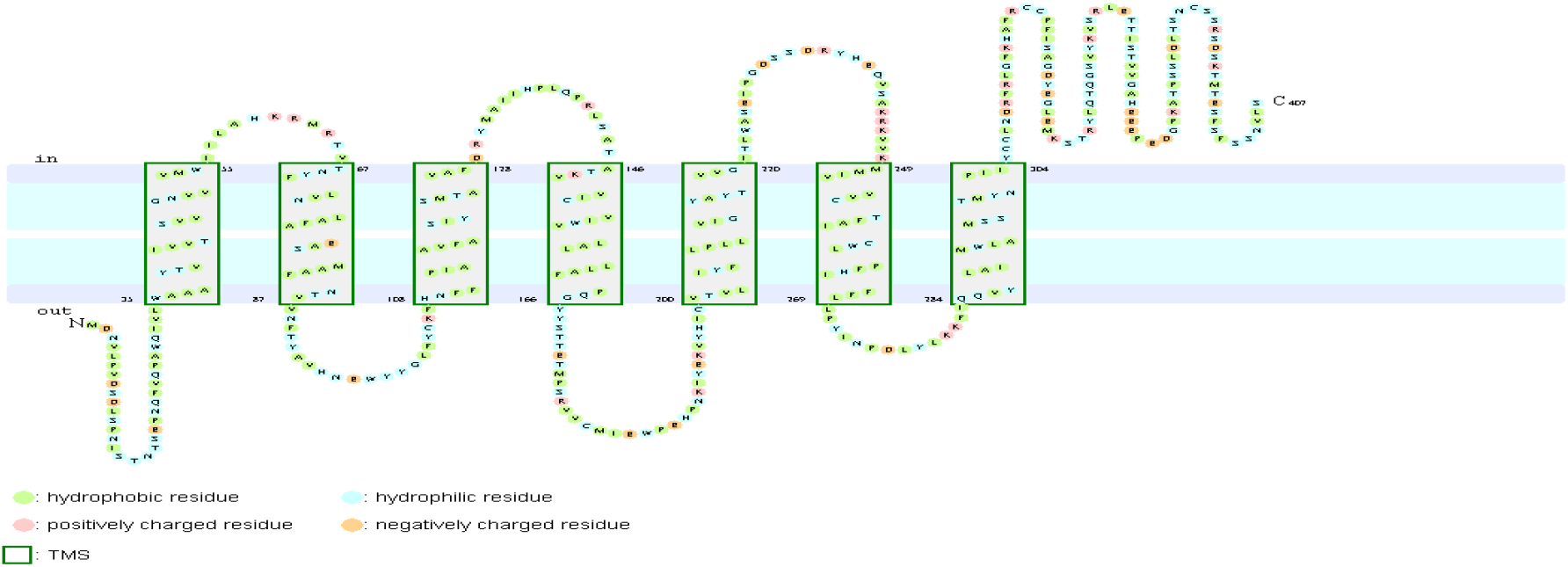
Transmembrane topology prediction of NK-1R by using ConPred II

NK-1R, stimulated directly, may cause vasodilation of feto-placental blood vessels[21]. In a study, SP has been found to be expressed in trophoblastic cell tissues, but not observed in blood vessels of fetus. mRNA of NK-1R has been reported in placental tissue[22].

Accordingly, the present study aimed to clarify the localization and expression for NK-1R in human retained products of conception. Role of NK-1R is not known in retained products of conception. Identification of the cause is challenging although, and there are no effective measures available for treatment[23].

## Methods

### Tissue Samples

The present study was approved by the Ethical Review Committee of The University of Lahore. Ten samples of POCs after different stages of spontaneous miscarriages were obtained from Gynaecology Department, The University of Lahore Teaching Hospital, Lahore. The ages of the females ranged from 20 to 40 years with mean age of 29 years. The gestational age ranged from 01 and 12 weeks and the mean weight of the fragments were 295 g.

### Immunohistochemical Staining for NK-1R

Human POC tissues were studied by immunohistochemistry to demonstrate the localization of NK-1R. Tissues of POCs were fixed in 10% formalin and then embedded in paraffin. 4-μm-thick serial sections were cut and processed for immunohistochemical detection of NK-1R. Antigen retrieval was done in 10 mM citrate buffer (pH 6.0), followed by peroxidase-blocking (Dako Cytomation A/S) on the sections and then heated at 100 °C for 60 min. NK-!R antibody (Abbott, 1:1000) was applied and incubated for 30 min at 37 °C. Then incubated in Universal secondary antibody (Roche Diagnostics KK) for 20 min at 37 °C. DAB Map detection kit (Roche Diagnostics KK) was used for visualization. Nuclei were counterstained using a hematoxylin counterstain reagent (Roche Diagnostics KK).

We studied representative samples of POCS. The evaluation of all slides were done by two independent pathologists. In each slide, high-power microscopic fields were evaluated using a 10, 20 and 40x magnification. The presence or absence of staining and the intensity of the immunoreactivity were noted, as well as the number and type of cells. Intensity of staining was observed as a brown staining. The localization of staining, whether or not the staining was localized in the nucleus, cytoplasm or in the plasma membrane was also observed. The results were recorded as positive when they showed cellular and/or plasma membrane staining ranging from moderate to strong in more than 10% of the cells. By consensus among the pathologists, the intensity of the immunoreactive cells was scored as follows: when less than 10% of the total cells were stained, the number of immunoreactive cells was considered low (+1), it was considered moderate when 10-40% were stained (+2) and high when more than 40% were stained (+3) [24]. The specimens were examined and photo-graphed at ×200 magnification utilizing a digital microscope camera (Olympus AX80 DP21; Olympus, Tokyo, Japan) interfaced with a computer. All protein levels were evaluated using the nuclear labeling index (%), recorded as the percentage of positively stained nuclei in 100 cells in selected area.

## Results

### NK-1R Protein Expression in products of conception Determined by Immunohistochemical Staining

Brains tissue was taken as a positive control for NK-1R and it showed intense staining of +3 in all the cells (Figure 2A, B). The expression of NK-1R protein was high in all the cases of POCs when evaluated in all the stages of miscarriages. NK-1R expression showed no notable differences among different cases of miscarriages irrespective of the age of females and the gestational age at which the event occurred. The NK-1R was widely distributed in the fetal membranes and placental tissues. The staining was high in epithelial cells of decidua, trophoblast of the fetal membranes and chorionic villi (cytotrophoblast and syncytiotrophoblast). NK-1R was expressed in all the cells of POCs, whether maternal or fetal, in their epithelial membranes and nuclei. We determined the immunohistochemical staining of NK-1R, which is expressed mainly in the brain (Figure 1F). NK.1R levels in products of conception at all the stages of miscarriage (Figure 1D) and G3 (Figure 1E) were higher than those in placental tissue after normal delivery (Figure 1B). Furthermore, we evaluated the nuclear labeling index of NK-1R using semi-serial sections (Figure 3). NK-1R was more strongly expressed in POCs. The NK-1R showed no difference in all the cases indicating that females who had spontaneous abortions have significantly higher levels of NK-1R.

**Figure 2.**
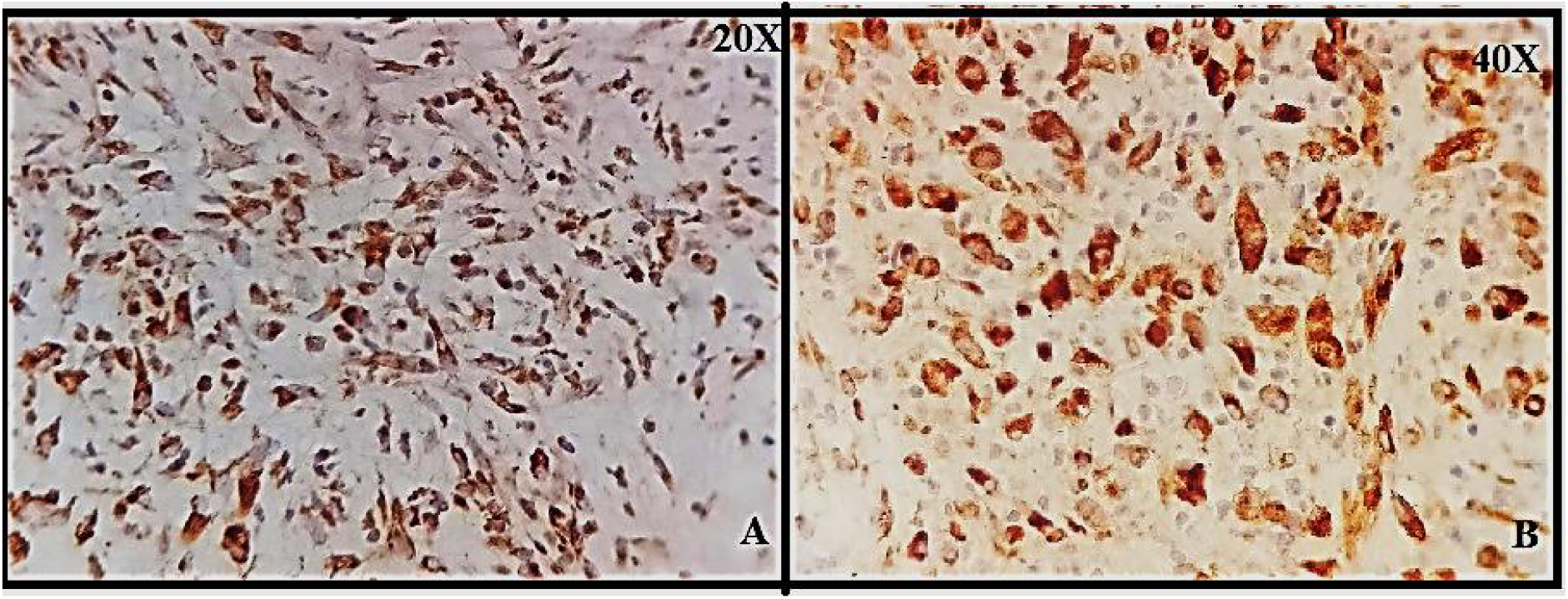
A.B: granular cytoplasmic positive staining in brain glial cells; positive control for NK-1R

**Figure 3.**
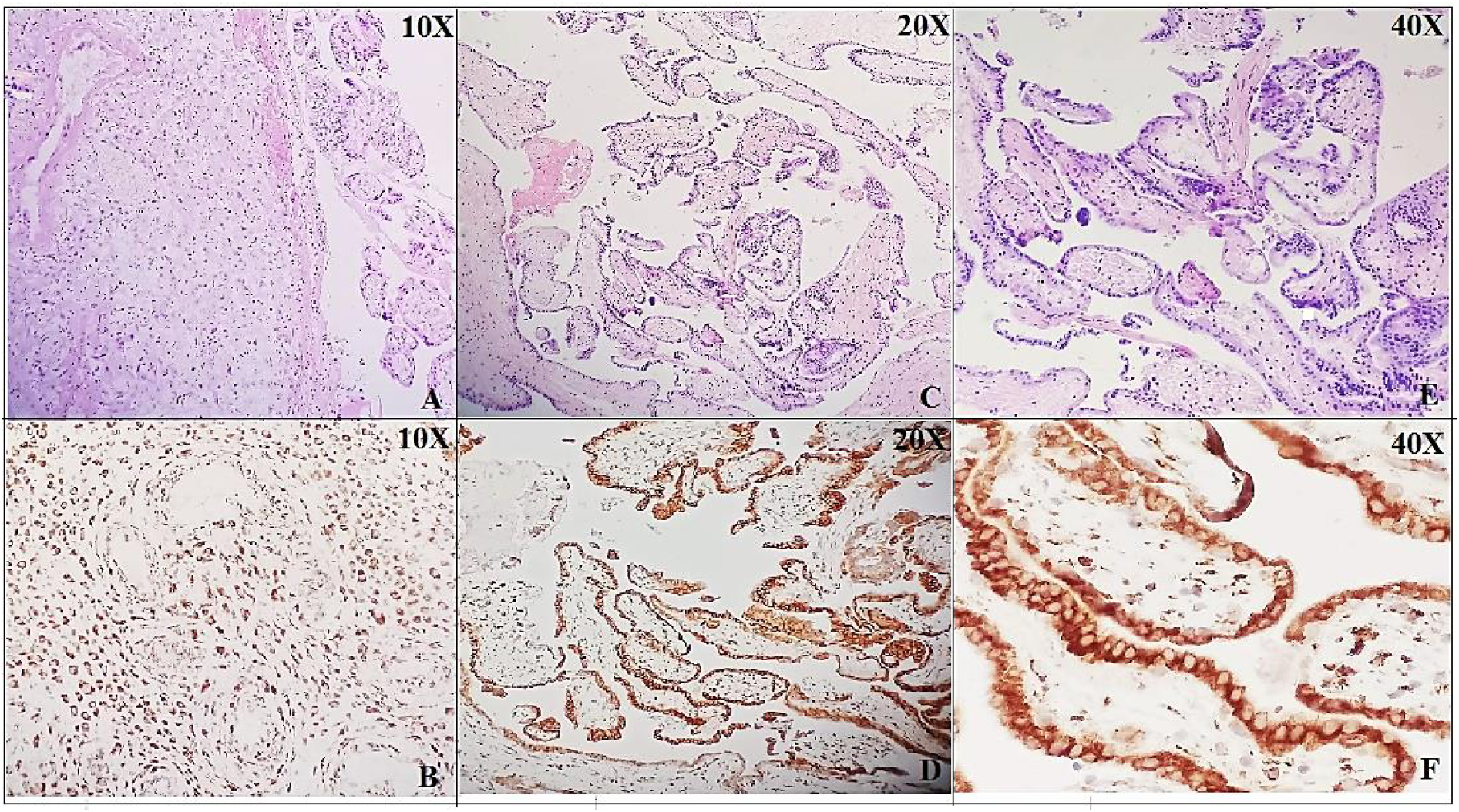
A: NK-1R immunohistochemical staining in products of conception showing intense staining and positive expression in all the cells at 10X (A,B), 20X (C,D) and 40X (E,F).

## Discussion

This study reports for the first time about the detailed expression of NK-1R in products of conception at an early gestational age in first trimester. Secondly, SP/NK-1 receptor system dysregulation may be involved in pathology of pregnancy, such as abortion [15,16], pre-eclampsia [25] and pre-term birth. We describe here for the first time the immunolocalization of NK-1R in human POCs and we provide evidence that NK-1R is expressed in the nucleus. All these observations suggest that the NK-1R and SP have a role in the pregnancy. In our view, the demonstration NK-1R in the in uterine products is associated with first trimester Miscarriages has an important functional implications. NK-1R has role in female reproduction. It has been known that mRNA for preprotachykinin-A which encodes SP is expressed in bovine corpus luteum (CL) of early developmental stage, CL with a retained oocyte shows that the muscular apparatus of the preovulatory follicle has a role in oocyte expulsion and follicle wall contraction was missing in the mutant group. This may suggest for luteinized unruptured follicle syndrome in humans [25].

There are several reports for the involvement of TKs in reproduction [26]. All the TKs and their receptors are found to be expressed in uterus of super-ovulated and unfertilized mice and may play a role in both male [27] and female reproductive system [28]. There is a significant up-regulation of NK-1R protein at full term fetus and newborn infant with a peak at day 1 and it downregulates at 8^th^ day, which indicates that NK-1R may be involved in the mechanisms modulating the processes during labour and after birth. SP-IR has an opposite correlation with NK-1R protein expression in pregnant and uterus after birth [29]. It correlates with our own previous study[30], in which it was proposed, based on our experimental work on sudden infant death syndrome, sudden fetal death victims and controls, [31] that SP expression is normally low in healthy fetuses but higher in neonates as compared to controls and if vice versa, it may lead to sudden death. Furthermore, it was proposed that the expression of SP is variable, depending on the developmental stage. It is lower in adults and if increased it may lead to death [30]. The possible reason is that SP is involved in cardiorespiratory control centers of the brain and has many important functions[32, 33]. It regulates and controls the sleep-wake cycle, respiratory rhythm as well. If there is any disturbance in its regulation, it may cause fatal outcomes including death. NK-1R is receptor of SP and all the functions of SP are only initiation, regulated and modulated, once SP binds with NK-1R. Here, in this study, we found an increased NK-1R expression which shows that if SP is increased in initial stage of pregnancy, it may cause spontaneous abortion or miscarriage; if at any fetal stage, it may initiate the respiratory mechanisms which is injurious as the lungs havnt yet started functioning and gaseous exchange is via feto-placental route. At full term, there is need to expel the fetus, there may be rise in SP expression, leading to birth of baby and same SP is required for respiration through lung after birth.. Munoz and colleagues observed the expression of SP and NK-1R in all the cells of placenta which is in line with our current and previous findings e.g. SP levels are raised at term fetus, near birth and soon after[22, 34]. We may speculate that SP is regulating the pregnancy and controlling the respiratory mechanisms, delivery and cardiorespiratory controls.

The role of SP in stress and anxiety is already established [35]. It is released more in such conditions as well as other nociceptive stimuli including pain[36]. Stress-induced abortions may induce a rise in decidual Tumor Necrosis Factor (TNF)-alpha and cause neurogenic inflammation. TNF-alpha is a possible stimulator for miscarriage [37] [38]. We already know that stress, pain anxiety may lead to spontaneous miscarriage or preterm delivery as well. It shows that the possible mechanism may be via SP/NK-1R system. A study reported similar findings, that SP expression was elevated in full term fetus and in newborm and may have played role in cervical ripening and labour [39].

Pregnancy is a unique physiological process. During the early developmental stages, an immune rejection caused by fetal antigens is inhibited by mother [25]. Immune system has a very important role in pregnancy. During implantation, the endometrium makes the maternal-fetal connection and recruits the T-regulatory cells to the site which is the local immune response [40]. If there is any insufficiency in this process, it may cause spontaneous abortion [41]. It may be speculated that increased immune reactions or hypersensitivity may cause immune rejection, leading to abortion. Immune dysfunction may have a significant role too.

Human immunoglobulin (IG) has been used widely for the treatment of abortion [42]. It may modulate the immune mechanisms [43] in such a way that it may tolerate the embryo. The immune dysfunction may be enhanced by IG + human chorionic gonadotropin (HCG) drug therapy and hence, improving the pregnancy outcomes. It needs to be further explored [44]. In a study, the outcomes were improved by 60% after the treatment. IG+HCG may be suggested to increase the rate of successful pregnancy by modulation of immune function [44]. The limitations of study is small sample size and absence of control group. It was not possible to have a control tissue but we have compared the results with our previous studies. Here, we suggest NK-1R antagonist in addition to the IG+HCG, to diagnose and treat spontaneous abortion.

